# Integrated Bioinformatics Analysis Reveals APEX1 as a Potential Biomarker for Septic Cardiomyopathy

**DOI:** 10.1101/2023.01.03.522553

**Authors:** Junxing Pu, Fan Gao, Ying He

**Affiliations:** School of Public Health, Nanchang University, Nanchang 330031, China; College of Life Science and Resources and Environment, Yichun University, Yichun 336000, China

## Abstract

**Background:** A severe threat to human health is septic cardiomyopathy (SCM), a condition with high morbidity and fatality rates throughout the world. However, effective treatment targets are still lacking. Therefore, it is necessary and urgent to find new therapeutic targets of SCM.

**Methods:** We obtained gene chip datasets GSE79962, GSE53007 and GSE13205 from the GEO database. After data normalization, GSE79962 was used as the training set and screened for differentially expressed genes (DEGs). Then, the module genes most related to SCM were identified via weighted gene co-expression network analysis (WGCNA). The potential target genes of SCM were obtained by intersection of DEGs and WGCNA module genes. We further performed Gene Ontology (GO) and Kyoto Encyclopedia of Genes and Genomes (KEGG) function and pathway enrichment analyses on these genes. In addition, potential biomarkers were screened using machine learning algorithms and receiver operating characteristic (ROC) curve analysis. Gene Set Enrichment Analysis (GSEA) was then used to explore the mechanisms underlying the involvement of potential biomarkers. Finally, we validated the obtained potential biomarkers in test sets (GSE53007 and GSE13205).

**Results:** A total of 879 DEGs were obtained by differential expression analysis. WGCNA generated 2939 module genes significantly associated with SCM. The intersection of the two results produced 479 potential target genes. Enrichment analysis showed that these genes were involved in the positive regulation of protein kinase A signaling, histone deacetylase activity and T cell receptor signaling pathway, etc. Then, the results of machine learning algorithm and ROC analysis revealed that NEIL3, APEX1, KCNJ14 and TKTL1 had good diagnostic efficacy. GSEA results showed that these genes involved in signaling pathways mainly enriched in base excision repair and glycosaminoglycan biosynthesis pathways, etc. Notably, APEX1 was significantly up-regulated in the SCM groups of the two test sets and the AUC (area under curve) > 0.85.

**Conclusions:** Our study identified NEIL3, APEX1, KCNJ14 and TKTL1 may play important roles in the pathogenesis of SCM through integrated bioinformatics analysis, and APEX1 may be a novel biomarker with great potential in the clinical diagnosis and treatment of SCM in the future.

## Introduction

Sepsis is a major threat to human health since it is one of the main causes of death in critically ill people around the world, and its incidence is rising each year [1][2]. It is noteworthy that the World Health Organization has identified sepsis as a major global public health problem [3]. In 2016, the European Society of Intensive Care Medicine and the American Society of Critical Care Medicine convened a working group of experts to reach a new consensus on the definition of sepsis: sepsis is a life-threatening organ dysfunction caused by a dysfunctional host response to infection [4].

SCM first described by Parker et al. in the 1980s, is an acute cardiac dysfunction caused by sepsis [5]. Although SCM can recover in the early stage of sepsis [6], as a common complication of sepsis, it is one of the main causes of morbidity and mortality in sepsis patients. Previous studies have shown that the incidence of sepsis with SCM is about 13.8 % −51.6%, and the mortality rate is as high as about 70%, which causes a serious threat to human life [6][9]. However, until now, it is still difficult to diagnose SCM without a gold standard [10]. In the past, most studies used the criterion that the left ventricular ejection fractions (LVEF) < 50% as the diagnostic index for SCM [11][13]. However, due to the fact that LVEF is largely dependent on loading conditions, it has been recognized that LVEF is an uncertain marker of intrinsic cardiac function [14][15]. Furthermore, unfortunately, because of the complex pathogenesis of SCM, there are no effective therapeutic agents for SCM at this stage. And the current treatment strategy is still focused on controlling the primary disease and avoiding the subsequent SCM [16]. Therefore, it is necessary and urgent to explore new effective biomarkers of SCM to help understand its pathogenesis and apply them to the early diagnosis and treatment of SCM in the future.

In this study, we used an integrated bioinformatics approach to find potential diagnostic biomarkers of SCM. Specifically, we first obtained three microarray datasets from the GEO database and divided them into a training set and two test sets. Data from the training set were normalized and then subjected to differential analysis to screen for DEGs. WGCNA was then performed to identify key module genes. Next, the potential target genes of SCM were obtained after taking the intersection of DEGs and module genes. Then, the functions and pathways involved in these potential target genes were investigated using GO and KEGG enrichment analyses. We further used machine learning algorithms to analyze these genes to find potential biomarkers. In addition, ROC and GSEA analyses were performed on the obtained potential biomarkers to evaluate their diagnostic efficacy for SCM and the possible mechanisms involved. Finally, the expression levels and diagnostic efficacy of the potential biomarkers were validated in two test sets, so as to obtain the potential diagnostic biomarkers of SCM. The workflow of this study is shown in Figure 1.

**Figure 1:**
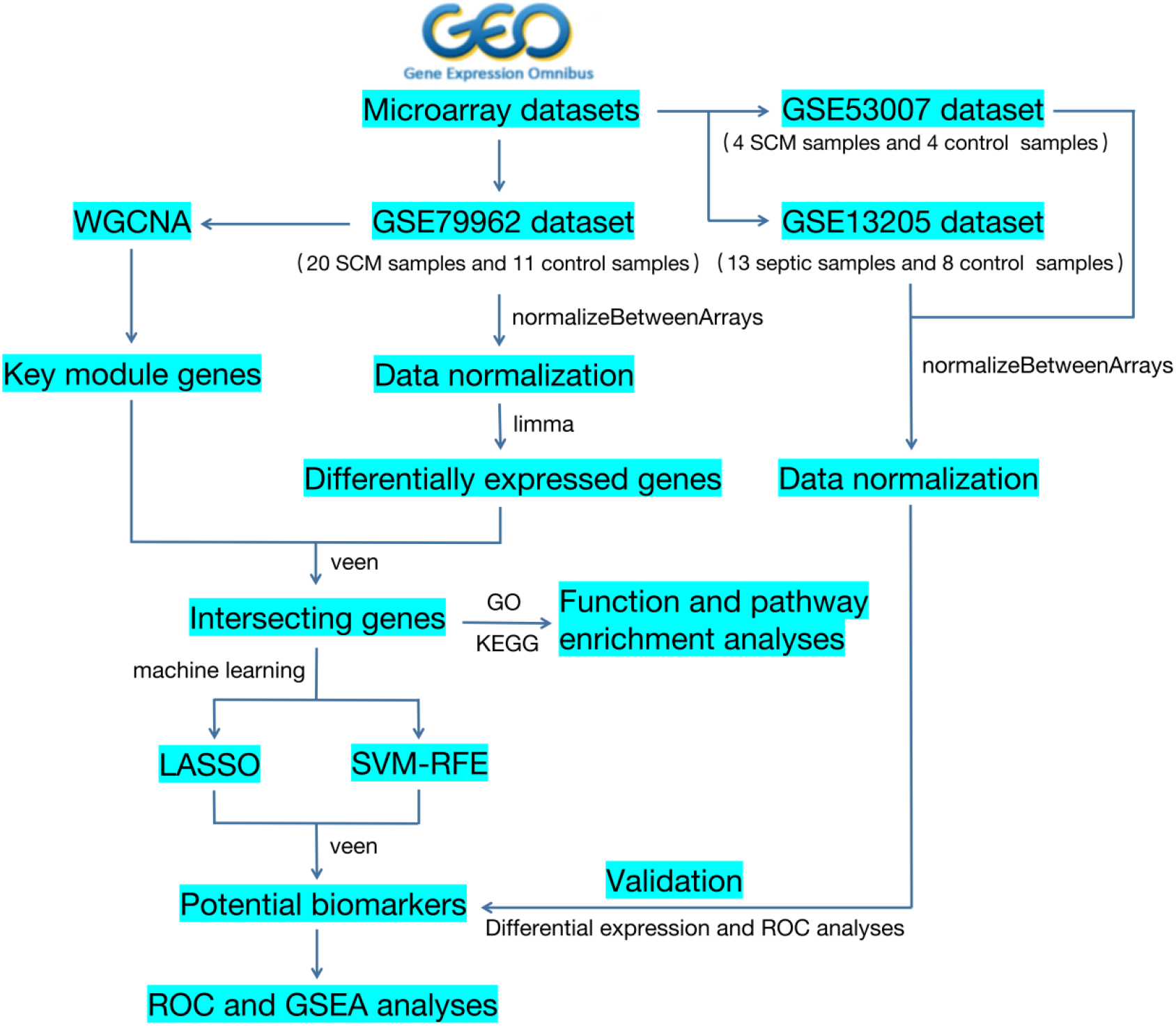
Workflow chart of this study.

## Materials and Methods

### Data Acquisition and Processing

The gene expression microarray datasets (GSE79962, GSE13205 and GSE53007) were downloaded from the GEO database (https://www.ncbi.nlm.nih.gov/geo/) [17]. The GSE79962 dataset based on the GPL6244 [HuGene-1_0-st] Affymetrix Human Gene 1.0 ST Array [transcript (gene) version] sequencing platform contains 20 heart tissue samples from SCM patients and 11 non-failing donor heart tissue samples. The GSE13205 dataset contains 13 muscle tissue samples from sepsis patients and 8 muscle tissue samples from control group. These muscle biopsy tissues were taken from the lateral portion of the lateral femoral muscle 10-20 cm from the knee of the corresponding study subjects. The GSE53007 dataset contains four septic mouse myocardial samples and four normal control myocardial samples sequenced on the Illumina MouseRef-8 v2.0 expression beadchip. For the case of multiple probes corresponding to the same gene name, the average value was taken for subsequent analysis. At the same time, we used the “normalizeBetweenArrays” method to correct the data to eliminate the batch effect as much as possible. In this study, the data set GSE79962 was used as the training set, while the test sets GSE13205 and GSE53007 were used to validate the potential biomarkers obtained from the training group at the cardiac and skeletal muscle levels, respectively.

### Screening for DEGs

After normalizing the data of the training set, the “limma” package in R software (version: 4.2.1) was used for difference analysis to obtain DEGs between the SCM group and the control group. The screening criteria for DEGs was adjusted *P* value (adj.*P*.Val) < 0.05 and |log2FC| > 0.4. The “pheatmap” and “ggplot2” packages in R software were used to display the heat map and volcano map of DEGs.

### Construction of Weighted Gene Co-expression Network

Based on the gene expression profile data of the training set, we used the “goodSamplesGenes” method to remove outliers and genes, and then used the “WGCNA” package [18] to construct a scale-free gene co-expression network. After the optimal soft threshold β was calculated by the “pickSoftThreshold” function, the weighted correlation adjacency matrix was constructed. Then, the adjacency matrix was transformed into a topological overlap matrix (TOM) and a hierarchical clustering and dynamic tree cutting algorithm was used to construct gene modules. The minimum number of genes within a module was set to 50 as well as merging similar modules with cut heights less than 0.25.

### Identification of Potential Target Genes

To determine the clinical significance of each gene module, we further analyzed the correlations between the module eigengenes (MEs) and clinical traits in each module. Then, associations between module expression patterns associated with sample traits were reflected by calculating gene significance (GS) and module membership (MM). We obtained the potential target genes in SCM by intersecting the important module genes obtained by WGCNA method with DEGs.

### Function and Pathway Enrichment Analyses

To further explore the functions and signaling pathways involved in potential target genes in SCM, we performed GO and KEGG enrichment analysis. GO analysis is used for annotation of gene functions, mainly biological process (BP), cellular component (CC) and molecular function (MF) [19]. KEGG is a commonly used online database for studying gene function and related high-level genomic functional information [20]. First, we used the R package “org.Hs.eg.db” for gene ID conversion and then the “clusterProfiler” package for enrichment analysis. *P* value < 0.05 was considered statistically significant.

### Potential Biomarkers Screening

Machine learning is a widely used technique in the field of artificial intelligence to analyze existing data to obtain patterns and use the patterns to predict unknown data [21]. In this study, two machine learning algorithms, LASSO logistic regression (“glmnet” package) and SVM-RFE (“e1071” package), were used to analyze potential target genes in SCM and the intersection of the obtained results were extracted. The intersecting genes were considered as the signature genes in SCM. Then, we used the R package “rms” to construct a nomogram model based on the signature genes. Finally, the diagnostic values of the signature genes and the nomogram model were evaluated in SCM samples and normal control samples using ROC curve analysis. The ROC curves were analyzed and visualized using the “pROC” package and “ggplot2”. If the AUC of the signature gene is > 0.8, it is considered to have excellent diagnostic efficacy and potential as a disease biomarker.

### Correlation and GSEA Analysis

We used the R package “corrplot” to correlate the expression of potential biomarkers in the training set. In addition, to further explore the KEGG signaling pathway involved in SCM, GSEA was performed based on the “clusterProfiler” software package for each of these potential biomarkers. *P* value < 0.05 was considered statistically significant.

### Validation of Potential Biomarkers

Based on the number of samples in the training set, we first analyzed the expression levels of potential biomarkers using a Student’s t-test. Then, after normalization in the test sets (GSE53007 and GSE13205 datasets), we performed difference analysis using the R package “limma” to verify whether the expression of potential biomarkers was consistent with that in the training sets. And *P* value < 0.05 was considered statistically significant. Finally, we also evaluated the diagnostic efficacy of them in the test sets.

## Results

### Identification of DEGs

As shown in Figures 2A and B, we first normalized the samples and then obtained 879 DEGs from the SCM and control groups. Among them, the statistically significant differentially expressed up-regulated and down-regulated genes were 411 and 468, respectively (Figure 2C). The expression of the top 30 up-regulated genes and the top 30 down-regulated genes with the most significant differences are shown in Figure 2D.

**Figure 2:**
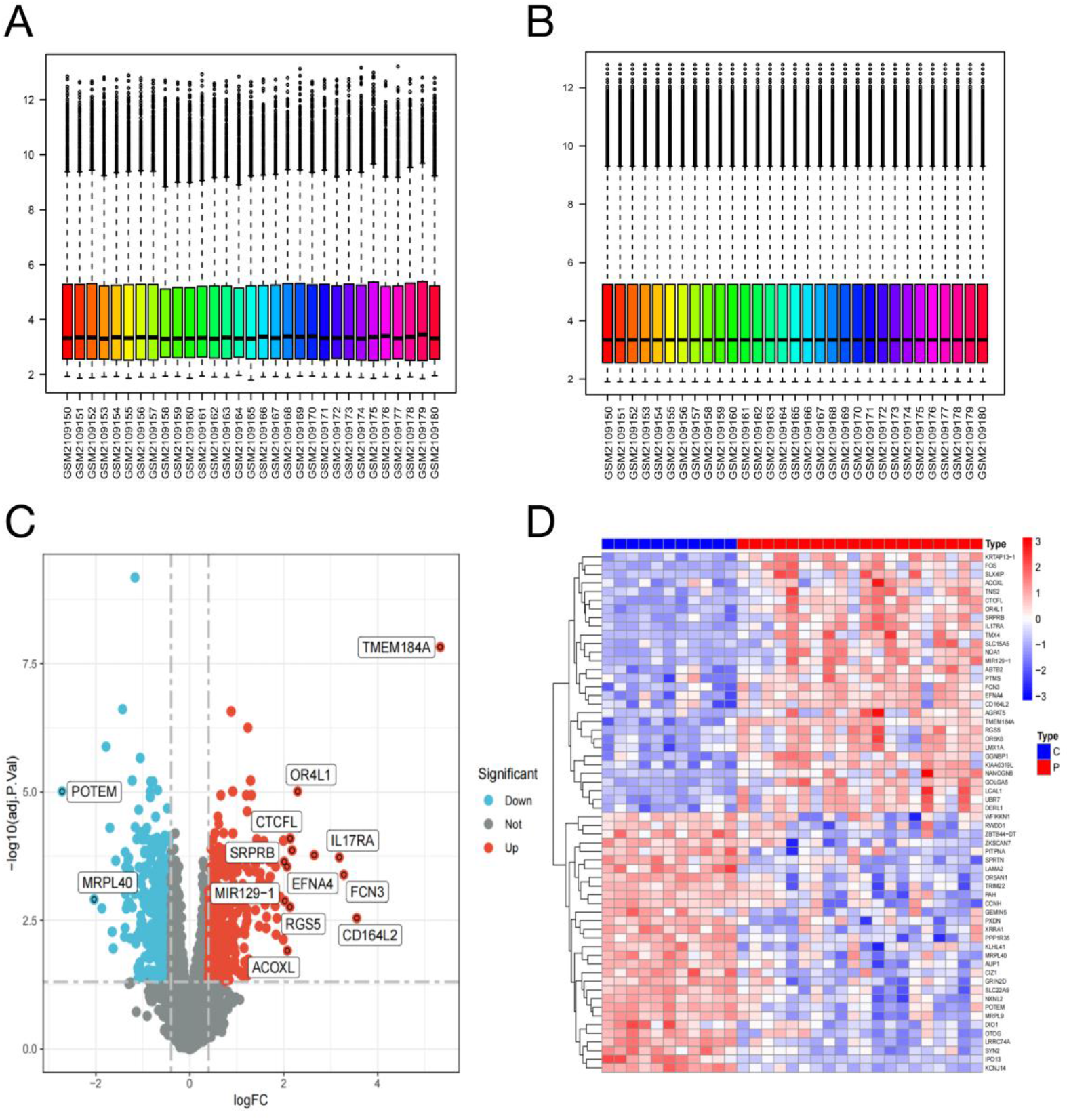
Identification of DEGs in the myocardium tissue of patients with SCM. (A) Training set data before normalization process. (B) Training set data after normalization process. (C) Volcano plot shows the expression of DEGs (adj.*P*.Val < 0.05 and |log2FC| > 0.4). Red and blue dots represent significant up- and down-regulation of genes in the SCM group, respectively. (D) Heatmap of the top 30 DEGs between SCM and control groups. Notes: C, control group; P, SCM group.

### WGCNA and Potential Target Gene Analyses

In this study, the best soft threshold β =5 corresponding to the scale-free fit index R^2^ of 0.9 for the first time was selected to construct the scale-free network (Figure 3A). The minimum number of genes within a module was set to 50. 18 modules were obtained after merging similar modules with cut height less than 0.25 (Figure 3B). Notably, the grey60 module was considered as the set of genes that could not be assigned to any module, so it was excluded first in the subsequent analysis. In the correlation analysis of module genes with clinical characteristics, magenta module, turquoise module, cyan module, lightyellow module, brown module and midnightblue module were significantly associated with SCM (Figure 3C). As shown in Figure 3D, the turquoise module was the most correlated with SCM in the GS-MM correlation analysis (cor = 0.79, *P* < 1e-200). Finally, the intersection of turquoise module genes with DEGs was taken to obtain 479 potential target genes, which were significantly related with SCM (Figure 3E).

**Figure 3:**
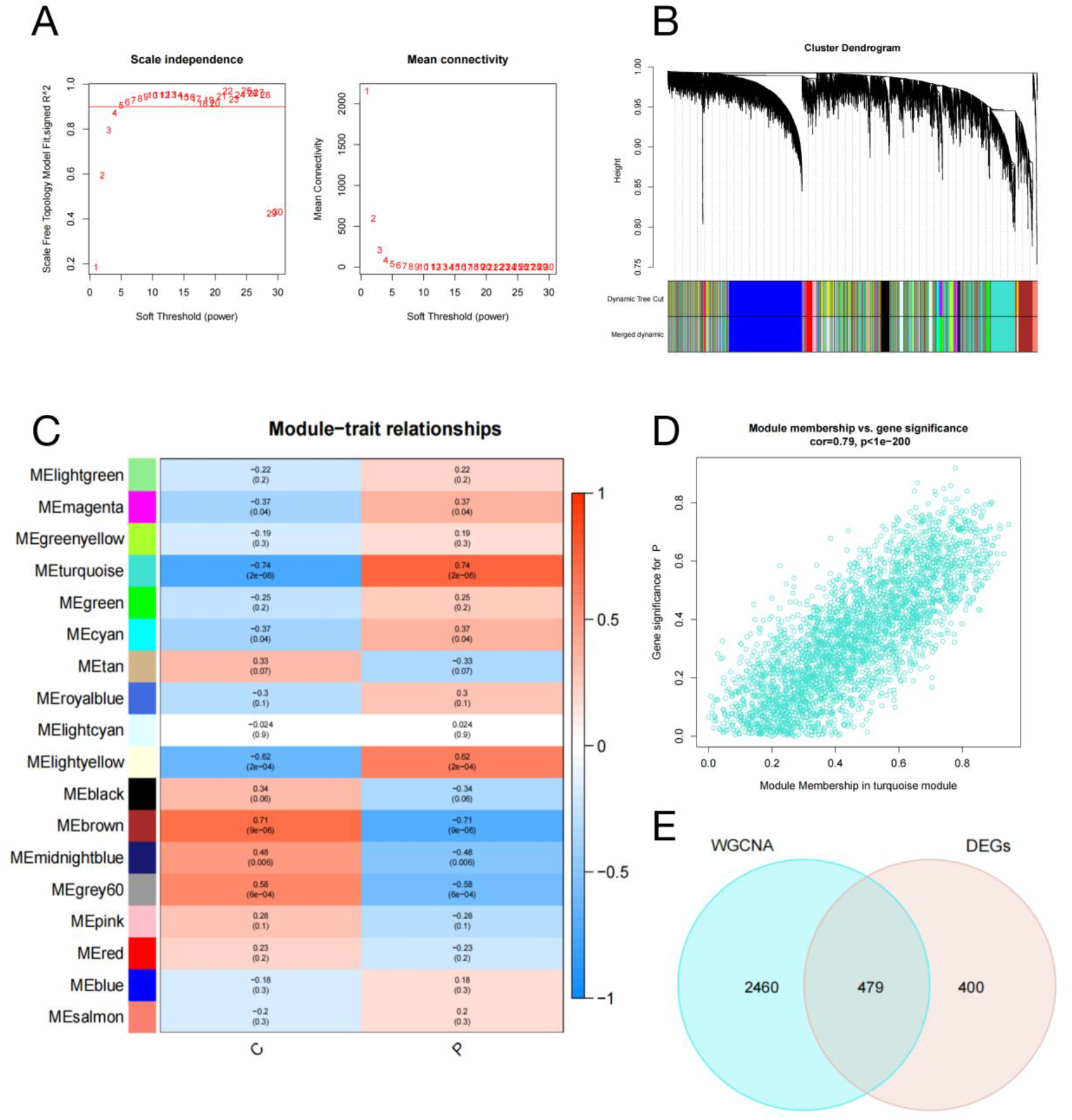
Screening of SCM-related key modules. (A) Network topology analysis for various soft threshold powers. Left: Scale-free fit index (y-axis) analysis for different soft thresholds (x-axis); Right: average connectivity (y-axis) analysis for different soft thresholds (x-axis). (B) Dendrogram of gene clustering based on topological overlap of dissimilarity. Each color represents a gene module. (C) Module-trait correlation. Each row corresponds to a module eigengene (ME) and each column to a clinical trait. Notes: C, control group; P, SCM group. (D) Scatter plot of gene significance vs. module membership for SCM in the turquoise module (cor = 0.79, *P* < 1e-200). (E) Intersection between WGCNA turquoise module genes and DEGs.

### GO and KEGG Enrichment Analyses

To study the functions of these 479 potential target genes and the pathways involved in them, GO and KEGG enrichment analyses were performed in our work. The results of GO analysis included 261 biological processes (BPs), 54 molecular functions (MFs) and 23 cellular components (CCs). We only showed the first 7 significantly enriched results, respectively (Figures 4A, B). Specifically, we found that BPs were mainly enriched in the positive regulation of protein kinase A signaling, positive regulation of protein serine/threonine kinase activity, substrate-dependent cell migration, regulation of ceramide biosynthetic process, extracellular matrix disassembly, regulation of sphingolipid biosynthetic and membrane lipid metabolic processes, etc. MFs were mainly associated with histone deacetylase activity, protein lysine deacetylase activity, carboxylic acid binding, DNA N-glycosylase activity, alpha-actinin binding, collagen binding and NAD-dependent histone deacetylase activity, etc. The CCs results showed that these potential target genes were mainly enriched in nuclear speck, cyclin-dependent protein kinase holoenzyme complex, sarcoplasmic reticulum membrane, ciliary base, rough endoplasmic reticulum membrane, clathrin vesicle coat and nuclear cyclin-dependent protein kinase holoenzyme complex. In addition, we obtained a total of six statistically significant results in the KEGG signaling pathway enrichment analysis, which were IL-17 signaling pathway, pentose phosphate pathway, phenylalanine metabolism, base excision repair, fructose and mannose metabolism and T cell receptor signaling pathway (Figures 4C, D).

**Figure 4:**
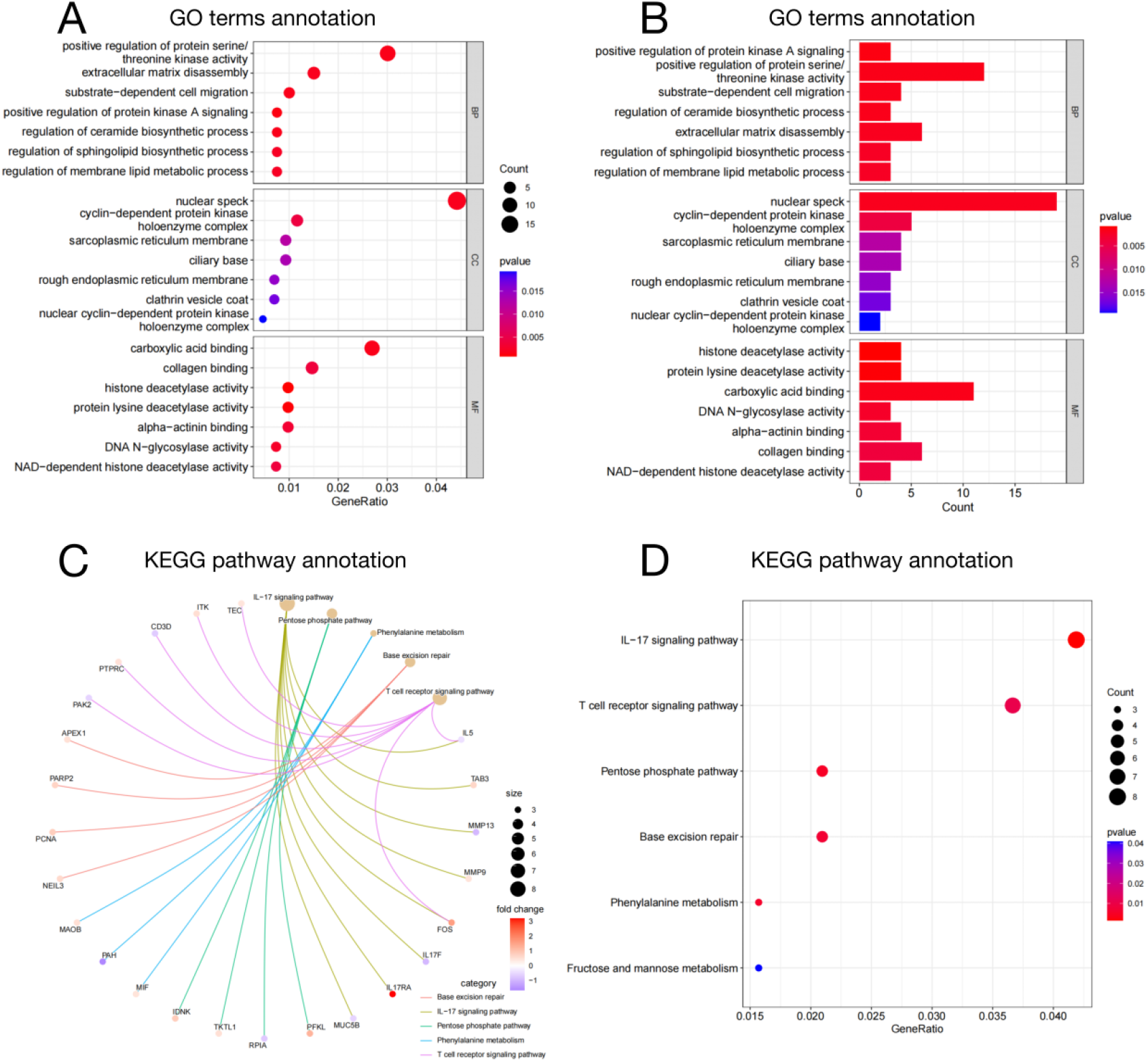
Function and pathway enrichment analyses of potential target genes in SCM. (A) Bubble plot of GO enrichment analysis showing top seven BPs, CCs and MFs. (B) Bar plot of GO enrichment analysis showing top seven BPs, CCs and MFs. (C) Cnet plot of KEGG enrichment analysis. (D) Bubble plot of KEGG enrichment analysis. Abbreviations: GO, Gene Ontology; KEGG, Kyoto Encyclopedia of Genes and Genomes; BP, biological process; CC, cellular component; MF, molecular function.

### Screening of Potential Biomarkers for SCM

We use two machine learning algorithms to further analyze these 479 target genes to find potential biomarkers related to SCM. LASSO regression analysis showed that the model composed of 14 candidate biomarkers could accurately predict SCM (Figure 5A, B). As shown in Figures 5C and D, in the SVM RFE analysis using quintupling cross validation, when the number of characteristic genes in the model was seven to ten, this model had the highest accuracy and the lowest error rate for predicting SCM. In order to make the model more stable, the first 10 characteristic genes were selected as biomarkers. Then, the results of the two machine learning algorithms were intersected to obtain a total of four potential biomarkers of SCM, which were NEIL3, APEX1, KCNJ14 and TKTL1 (Figure 5E).

**Figure 5:**
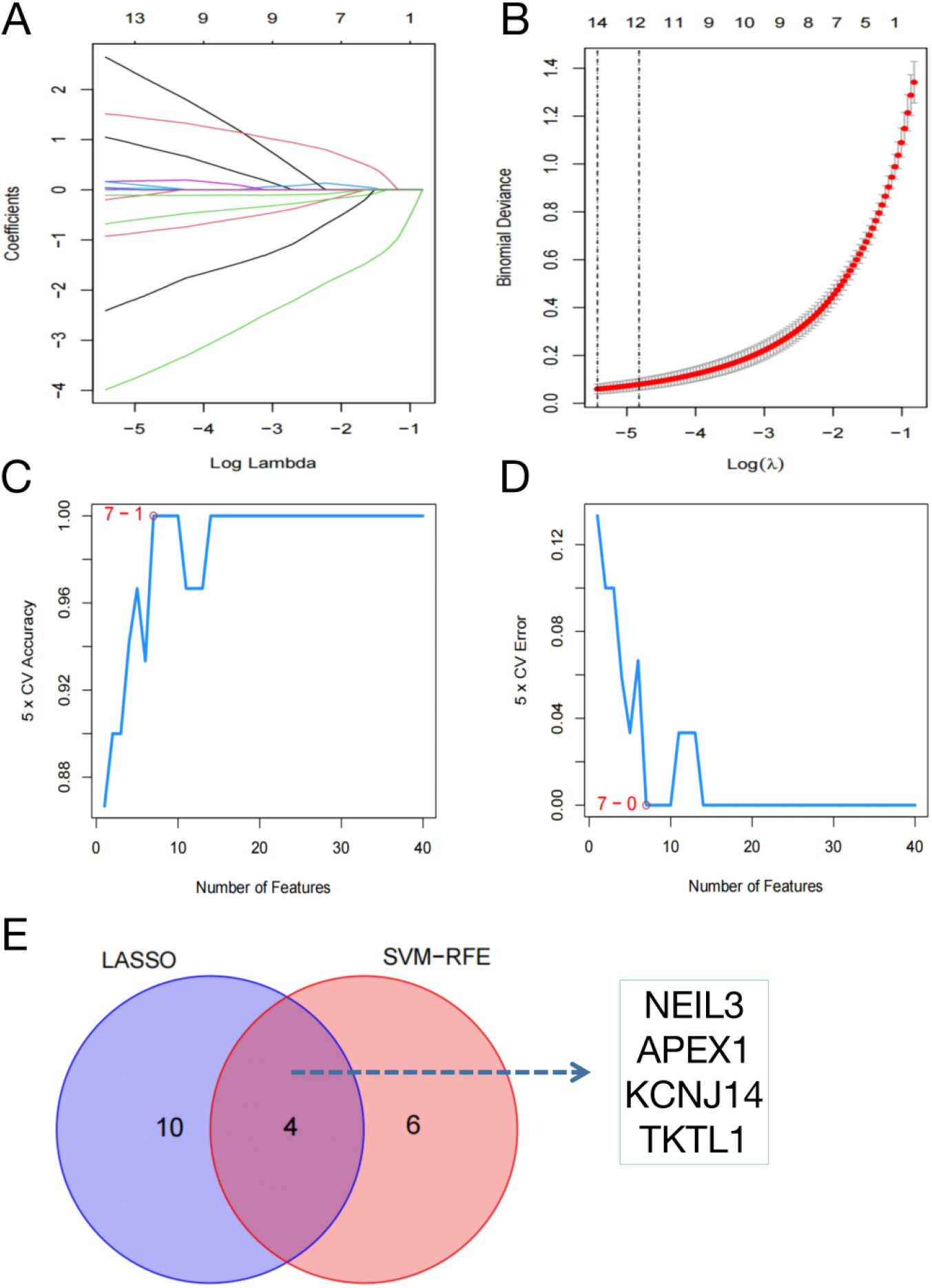
Exploration of potential biomarkers in SCM based on machine learning algorithms. (A) (B) LASSO regression model composed of 14 genes. (C) (D) The 5-fold cross-validation curve in the SVM-RFE model shows the highest accuracy and lowest error rate when the number of feature genes are seven to ten. (E) Four potential biomarkers in SCM (NEIL3, APEX1, KCNJ14 and TKTL1).

Furthermore, our study used these four potential biomarkers to construct a diagnostic prediction model. And the results of the nomogram showed that their expression contributed to the clinical diagnosis of SCM. We scored these potential biomarkers according to their expression to assess the likelihood of developing SCM (Figure 6A). Finally, ROC analysis was used to evaluate the diagnostic efficacy of these potential biomarkers and the nomogram model. As shown in Figures 6B-F, the AUCs of NEIL3, APEX1, KCNJ14, TKTL1, and the model were all > 0.95, which indicated that NEIL3, APEX1, KCNJ14, TKTL1 had the potential to be diagnostic biomarkers for SCM.

**Figure 6:**
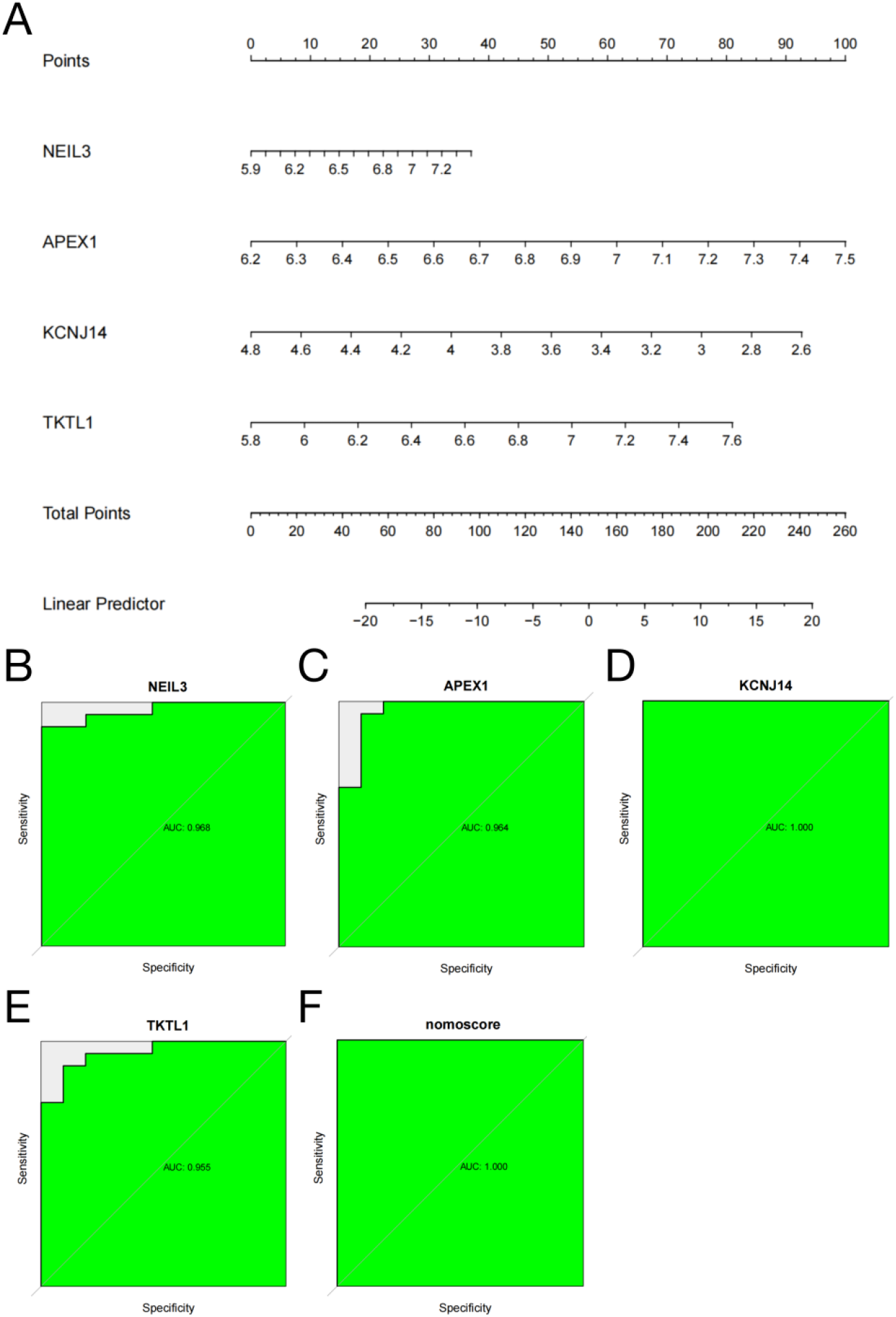
A diagnostic prediction model for SCM based on four potential biomarkers and evaluation of their diagnostic efficacy. (A) Nomogram. We assigned scores to each factor at different value levels according to the contribution of each factor to the outcome variable in the model, and then summed the scores to obtain the total score and to predict the risk of SCM in patients. (B) ROC curve showing the diagnostic efficacy of NEIL3 (AUC: 0.986). (C) ROC curve showing the diagnostic efficacy of APEX1 (AUC: 0.964). (D) ROC curve showing the diagnostic efficacy of KCNJ14 (AUC: 1.000). (E) ROC curve showing the diagnostic efficacy of TKTL1 (AUC: 0.955). (F) ROC curve showing the diagnostic efficacy of nomoscores (AUC: 1.000).

### GSEA and Correlation Analysis of Potential Biomarkers

To explore the possible mechanisms involved in these four potential biomarkers in SCM, we performed GSEA analysis to predict their KEGG downstream pathways, respectively. As shown in Figures 7A-D, the signaling pathways in the high NEIL3, APEX1, KCNJ14 or TKTL1 expression groups were mainly enriched in aminoacyl-tRNA biosynthesis, base excision repair, one carbon pool by folate, regulation of lipolysis in adipocytes, retinol metabolism, ABC transporters and drug metabolism-cytochrome P450. The signaling pathways in the low NEIL3, APEX1, KCNJ14 or TKTL1 expression groups were mainly enriched in chronic myeloid leukemia, glycosaminoglycan degradation, mucin type O-glycan biosynthesis, apoptosis-multiple species, small cell lung cancerand Toll-like receptor signaling pathway. The correlation result showed that KCNJ14 was significantly negatively correlated with the expression levels of NEIL3, APEX1 and TKTL1 in myocardial tissues of SCM patients, while NEIL3 was significantly and positively correlated with both APEX1 and TKTL1 (Figures 7E, F).

**Figure 7:**
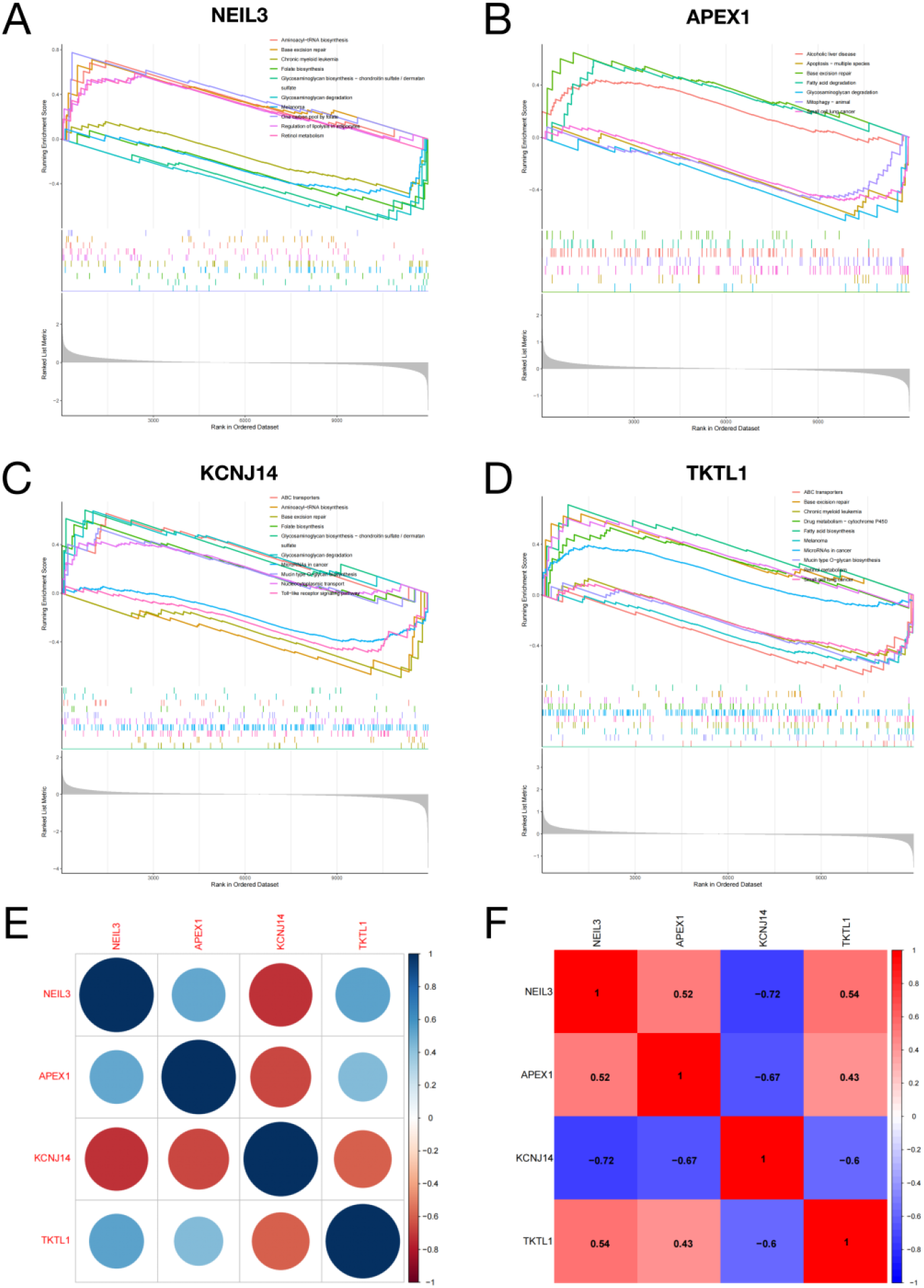
GSEA pathway enrichment analysis of potential biomarkers and their correlation analysis. (A-D) GSEA signal pathway enrichment analysis of NEIL3, APEX1, KCNJ14 and TKTL1. (E) (F) Correlation analysis.

### Expression Characteristics of Potential Biomarkers with External Validation

Subsequently, we examined the expression characteristics of NEIL3, APEX1, KCNJ14 and TKTL1 in the training set, and the results showed that NEIL3, APEX1 and TKTL1 were significantly upregulated in SCM (Figures 8A, B, D), while KCNJ14 was significantly downregulated (Figure 8C), which was the same as the results obtained in the correlation analysis. Since the expression information of NEIL3, KCNJ14 and TKTL1 was not available in the myocardial test group GSE53007, we mainly focused on assessing the expression characteristic and disease diagnostic efficacy of APEX1 in the test sets GSE53007 and GSE13205 datasets. As shown in Figures 8E, G, APEX1 expression was significantly upregulated in both myocardial tissue of septic mice and skeletal muscle tissue of septic patients in the test group, consistent with its results in the training set. In terms of ROC analysis, we found that APEX1 also had good disease diagnostic efficacy in both test sets (GSE53007 dataset: AUC: 1.000; 95% CI: 1.000-1.000, Figure 8F; GSE13205 dataset: AUC: 0.894; 95% CI: 0.712-1.000, Figure 8H). From the above analysis, we suggested that NEIL3, APEX1, KCNJ14 and TKTL1 might play an important role in SCM and APEX1 had a high potential to become a novel diagnostic biomarker for SCM.

**Figure 8:**
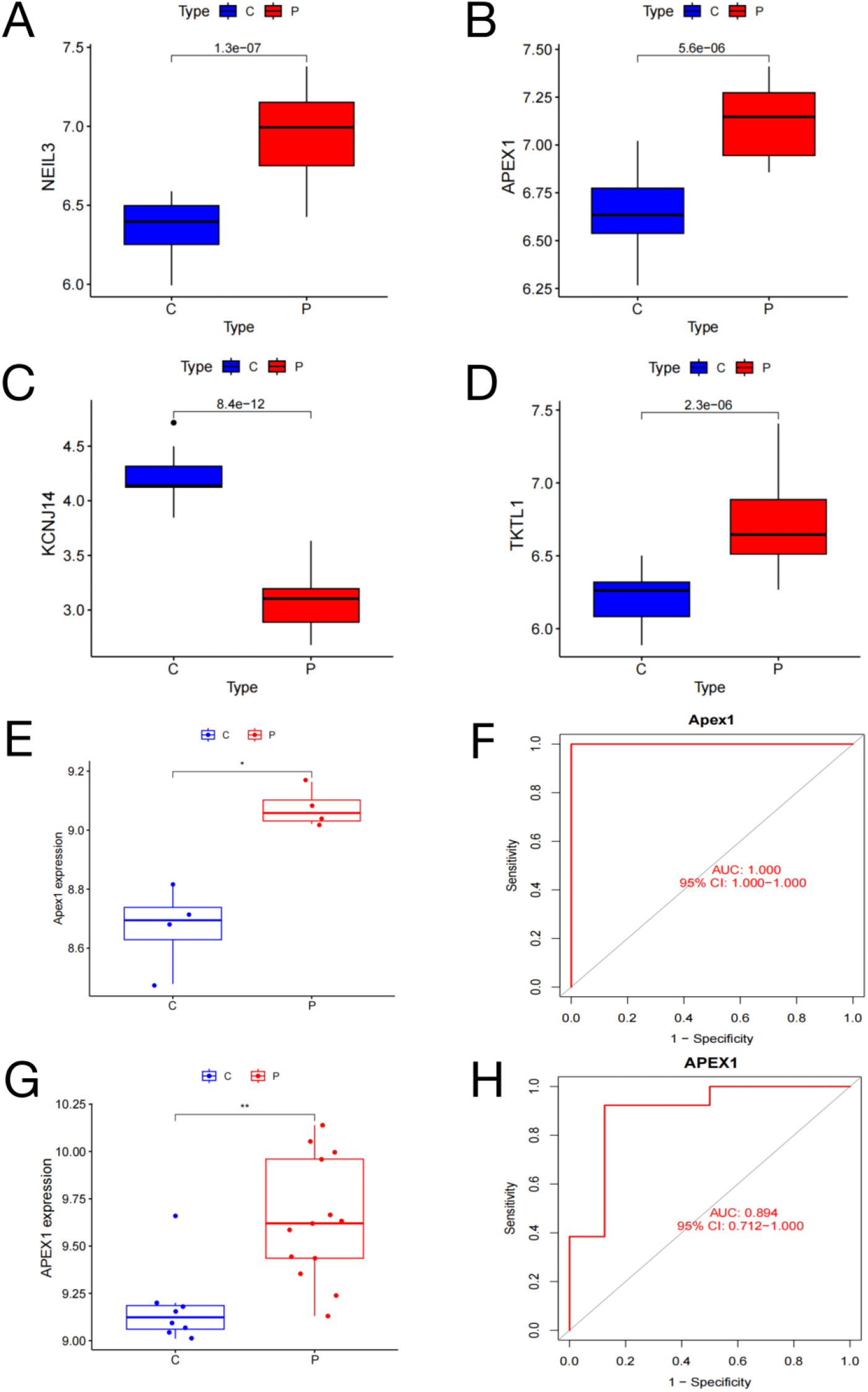
Analysis of the expression characteristics of potential biomarkers in the training set and validation of APEX1 expression level and diagnostic efficacy in two test sets. (A-D) NEIL3, APEX1 and TKTL1 expression were all significantly upregulated in SCM, while KCNJ14 expression was significantly downregulated. (E) (F) Expression and ROC evaluation of APEX1 in the GSE53007 dataset (expression significant upregulation and AUC=1.000). (G) (H) Expression and ROC evaluation of APEX1 in the GSE13205 dataset (expression significant upregulation and AUC=0.894). Notes: C, control group; P, SCM group. *, *P* < 0.05; **, *P* < 0.01.

## Discussion

Sepsis is a systemic inflammatory response syndrome caused by infection [22], and it is also one of the main causes of death in intensive care unit patients [23]. The mortality rate is as high as 41.9%, which seriously threatens human life and health [24]. SCM is an acute cardiac dysfunction subsequent to sepsis. Although it can recover in the early stage of sepsis, it is still one of the main causes of death in patients with sepsis [25]. The pathogenesis of SCM is very complex and involves the interaction between multiple hosts and pathogens, which has not been systematically elucidated [26]. Due to the complexity of the pathogenesis and the lack of specific clinical manifestations, the early diagnosis of SCM is still difficult [10]. Studies have shown that it is necessary to combine clinical, electrocardiographic, hemodynamic parameters and echocardiography and other methods to fully diagnose SCM [27]. Previous studies have identified novel biomarkers in SCM by differential analysis and protein-protein interaction network analysis [28]. For the first time, our study combined difference analysis, WGCNA and machine learning algorithms to screen potential biomarkers of SCM. We also performed function analysis of the obtained biomarkers and constructed a nomogram diagnostic model. Two external datasets were used to validate the results from different perspectives (myocardial tissue and skeletal muscle tissue) to increase credibility. Our results suggested that these biomarkers have great potential in the diagnosis of SCM and may be helpful for future research on clinical therapeutic targets.

In this study, the data were first normalized, and then 879 DEGs were screened from the myocardial tissue of SCM patients by differential analysis. Subsequently, 18 co-expressed gene modules were obtained by WGCNA. Among these gene modules, magenta module, turquoise module, cyan module, lightyellow module, brown module and midnightblue module were significantly correlated with SCM. GS-MM correlation analysis showed that turquoise module had the strongest correlation with SCM (cor = 0.79, *P* < 1e-200). The 2939 genes within the turquoise module were intersected with DEGs to obtain 479 potential target genes for SCM. GO function analysis showed that their BPs mainly enriched in positive regulation of protein serine/threonine kinase activity, substrate-dependent cell migration, extracellular matrix disassembly, and membrane lipid metabolism. MFs were significantly correlated with histone deacetylase activity, protein lysine deacetylase activity, DNA N-glycosylase activity, alpha-actinin binding, and NAD-dependent histone deacetylase activity. CCs results mainly enriched in nuclear speck, cyclin-dependent protein kinase holoenzyme complex and rough endoplasmic reticulum membrane. In addition, our enriched results in KEGG signaling pathway analysis included IL-17 signaling pathway, pentose phosphate pathway, phenylalanine metabolism, base excision repair, fructose and mannose metabolism, and T cell receptor signaling pathway.

In addition to cell phagocytosis and cytokine production, substrate-dependent cell migration is one of the host responses to invading pathogens during inflammation [29]. Previous studies have shown that SCM is closely related to inflammation, oxidative stress and mitochondrial damage [30]. The experimental results of Zhong et al. showed that endoplasmic reticulum stress was also related to the progress of SCM [31]. In terms of signaling pathways, Li et al. revealed that the IL-17 pathway played an important role in the sepsis mouse model and they demonstrated that blocking IL-17A with neutralizing antibodies could alleviate sepsis-induced intestinal motility disorders [32]. In recent years, studies have shown that cell metabolism reprogramming plays an important role in the activation of the immune system and excessive inflammatory response [33][34]. Li et al. used metabolomics analysis to observe the metabolites of mouse bone marrow-derived macrophages induced by lipopolysaccharide (LPS), they found that these metabolites were mainly enriched in glucose metabolism pathways such as fructose and mannose, and pointed out that glucose metabolism might play an important role in sepsis inflammation [29]. These studies were highly correlated with the results of our function and pathway analysis, which could provide strong support for our results.

Subsequently, we screened the target genes of SCM by LASSO regression and SVM-RFE machine learning algorithms, and then combined the nomogram diagnostic model and ROC analysis to identify whether NEIL3, APEX1, KCNJ14 and TKTL1 in myocardial tissue can be used as potential diagnostic biomarkers of SCM. GSEA analysis was performed on them, and the enriched pathways mainly included base excision repair (BER) and glycosaminoglycan biosynthesis pathways, which were consistent with the results of our previous KEGG analysis of DEGs. In addition, correlation analysis showed that the expression of KCNJ14 was significantly negatively correlated with the expression of NEIL3, APEX1 and TKTL1 in the myocardial tissue of SCM patients, while the expression levels of NEIL3, APEX1 and TKTL1 were positively correlated with each other.

Nei like DNA glycosylase 3 (NEIL3) is a DNA glycosylase belonging to the bacterial Fpg/Nei family, and its homologues include NEIL1 and NEIL2 [35]. These DNA glycosylases can initiate the BER pathway by recognizing and cutting bases damaged by reactive oxygen species (ROS), thereby protecting cells from oxidative DNA damage [36] Furthermore, a previous study has shown that DNA glycosylases also play a key role in protecting telomere integrity [37]. Moreover, Hewitt et al. reported that telomere tended to be severer damaged and was shorter than other regions of the genome when it was under oxidative stress [38] [39]. These damages might cause telomere dysfunction. A recent study showed that the shorter telomere length in peripheral white blood cells was associated with higher mortality in critically ill patients, especially in sepsis [40]. Notably, Wang et al. demonstrated the ability of NEIL3 glycosylase to repair oxidative damage in telomeric DNA [41]. Although there is no direct evidence on the specific mechanism of NEIL3 glycosylase in sepsis or SCM, the results of the above-mentioned studies can help us to understand the possible role of glycosylase in countering the damage to the genome caused by oxidative stress in sepsis by initiating the BER pathway.

Apurinic/apyrimidinic endodeoxyribonuclease 1 (APEX1 or APE1), also known as redox effector factor 1 (Ref-1), is a multifunctional nucleic acid endonuclease that plays an important role in DNA damage repair and intracellular redox regulation [42]. Similar to NEIL3 glycosylase, APEX1 is also involved in the BER process. The difference is that APEX1 mainly cleaves apurinic/apyrimidinic (AP) sites generated by DNA glycosylases after excision of damaged bases, thus contributing to DNA damage repair [43]. APEX1’s role in the reductive regulation of antioxidant stress is mainly reflected in attenuating the inflammatory response. Park et al. showed that TNF-α-stimulated secretion of APEX1 by endothelial cells could play an anti-inflammatory role by disrupting TNFR1 inflammatory signaling [44]. In addition, Joo et al. demonstrated that APEX1 played an important anti-inflammatory role in LPS-induced sepsis mice and they suggested that secreted APEX1 might have a therapeutic effect on systemic inflammation [45].

Potassium inwardly rectifying channel subfamily J member 14 (KCNJ14), also known as inwardly rectifying K^+^ 2.4 (Kir2.4), is a member of the Kir2 subfamily of inwardly rectifying K^+^ channels [46]. The subunits of the Kir2 family (including Kir2.1, Kir2.2, Kir2.3, and Kir2.4) play important roles in excitability of various tissues such as the central nervous system and heart (heart rate regulation, the neurotransmitter release, and involvement in immune regulation) [47][48]. Previous studies have reported that ROS-induced oxidative stress interferes with the gating mechanism of K^+^ channels and may cause cardiac arrhythmias, T-lymphocyte apoptosis, and phagocytosis [49]. Yu et al. showed that Kir2.1-mediated plasma membrane potential in macrophages was a key regulator of the inflammatory response promoted by metabolic reprogramming and they found that pharmacological targeting of Kir2.1 attenuated inflammation in LPS-induced sepsis mice [50]. In addition, Li et al. suggested that KCNJ14 could be an independent prognostic risk factor for colorectal cancer patients and has the potential to be a biological target for colorectal cancer therapy [51]. However, the mechanism of action of KCNJ14 in sepsis or SCM is unclear and needs to be further investigated.

Transketolase like 1 (TKTL1) is a gene encoding a transketolase-like 1 protein that plays a catalytic role in the non-oxidative pathway of the pentose phosphate pathway (PPP) [52]. Current reports on TKTL1 mainly focused on tumors. For example, a study by Langbein et al. found that TKTL1 was significantly overexpressed in pulmonary metastatic colon cancer and that tumor metastasis was significantly inhibited when TKTL1 enzyme activity was inhibited with drugs [53]. Subsequent studies have also highlighted that overexpression of TKTL1 promoted cell proliferation and tumor growth as well as downregulation of TKTL1 expression inhibits the proliferation of various cancer cells [54][56]. It has been reported that TKTL1 overexpression could upregulate the expression of lactate dehydrogenase 5 (LDH5), which promoted the glycolytic process and produces large amounts of lactic acid, ultimately leading to apoptosis of normal cells surrounding the tumor and promoting tumor cell metastasis [57]. Notably, lactate is a typical biomarker of poor prognosis in patients with critical illnesses (especially sepsis), and its level is significantly correlated with patient morbidity and mortality [58]. Based on the above studies, we onclude that TKTL1 overexpression may be involved in the pathogenesis of SCM by promoting the production of large amounts of lactate. However, it still needs to be demonstrated by further experiments.

In this study, we performed an examination of APEX1 expression level and diagnostic efficacy in the test sets (GSE53007 and GSE13205) and found that APEX1 was highly upregulated not only in myocardial tissues of SCM but also in skeletal muscle tissues of sepsis. Moreover, the results of ROC analysis in both tested sets showed that APEX1 had excellent diagnostic efficacy for disease. Therefore, we suggest that NEIL3, APEX1, KCNJ14 and TKTL1 may play an important role in the pathogenesis of SCM and APEX1 has a great potential to become a novel diagnostic biomarker for SCM.

However, there are some limitations in our study. Firstly, the available transcriptomic datasets of SCM myocardial tissue are very limited. In this study, we only used one training set and two test sets. Second, the potential biomarkers we obtained still need more support from literature. Finally, these potential biomarkers need to be validated at the cellular and animal levels.

## Conclusions

In summary, through integrated bioinformatics analysis, we believe that NEIL3, APEX1, KCNJ14 and TKTL1 may play an important role in the occurrence and development of SCM. Validated by external datasets, we suggest that APEX1 has great potential to become a novel diagnostic biomarker and therapeutic target for SCM. The pathogenesis of SCM is very complex, which brings a great challenge to clinical diagnosis and treatment. Our study provides a new target for the early diagnosis and clinical treatment of SCM, but their specific mechanism in the pathogenesis of SCM still need to be further investigated in the future.

